# SARS-CoV-2 variant with higher affinity to ACE2 shows reduced sera neutralization susceptibility

**DOI:** 10.1101/2021.03.04.433887

**Authors:** Monique Vogel, Xinyue Chang, Gilles Sousa Augusto, Mona O. Mohsen, Daniel E. Speiser, Martin F. Bachmann

**Affiliations:** Department of Immunology, University clinic of Rheumatology and Immunology, Inselspital, Bern, Switzerland; Department of BioMedical Research, University of Bern, Bern, Switzerland; Nuffield Department of Medicine, Centre for Cellular and Molecular Physiology (CCMP), The Jenner Institute, University of Oxford, Oxford, UK; International Immunology Centre, Anhui Agricultural University, Hefei, China

**Keywords:** SARS-CoV-2, RBD, affinity, neutralization, vaccine, biolayer interferometry

## Abstract

**Background:** Several new variants of SARS-CoV-2 have emerged since fall 2020 which have multiple mutations in the receptor binding domain (RBD) of the spike protein.

**Objective:** We aimed to assess how mutations in the SARS-CoV-2 RBD affect receptor affinity to angiotensin-converting enzyme 2 (ACE2) and neutralization by anti-RBD serum antibodies.

**Methods:** We produced a SARS-CoV-2 RBD mutant (RBDmut) with key mutations (E484K, K417N, N501Y) from the newly emerged Brazilian variant. Using Biolayer Interferometry, we analyzed the binding of this mutant to ACE2, and the susceptibility to neutralization by sera from vaccinated mice and COVID-19 convalescent patients.

**Results:** Kinetic profiles showed increased RBDmut - ACE2 affinity compared to RBDwt, and binding of vaccine-elicited or convalescent sera was significantly reduced. Likewise, both sera types showed significantly reduced ability to block RBDmut - ACE2 binding indicating that antibodies induced by RBDwt have reduced capability to neutralize mutant virus.

**Conclusion:** Our physiochemical data show enhanced infectivity and reduced neutralization by polyclonal antibodies of the Brazilian variant of SARS-CoV-2.

**Capsule summary:** SARS-CoV-2 variant with Brazilian RBD mutations shows increased ACE2 affinity and reduced susceptibility to blockage by vaccine-elicited and convalescent sera.

## Introduction

Although no correlate of protection from SARS-CoV-2 has yet been confirmed, it is likely that this will apply to serum neutralization activity as it is a common correlate of protection against viral infection following natural infection or vaccination^1, 2^. Neutralizing antibody responses against SARS-CoV-2 are primarily directed against the receptor binding domain (RBD) of the spike protein. Within the RBD, the receptor binding motif (RBM) is the most important site, as it directly interacts with ACE2. The affinity of spike for ACE2 is 4-fold increased between SARS-CoV-1 and SARS-CoV-2, likely contributing to the increased infectivity and transmission of the latter^3, 4^. Likewise, antibodies against SARS-CoV-2 did not show cross-reactivity with RBD of SARS-CoV-1 indicating limited serum cross-neutralization and notable differences in the antigenicity between the closely related SARS-CoV-1 and SARS-CoV-2^5^. Recently, virus variants from the United Kingdom (N501Y, 60/70-deletions, A570D, D614G, P681H), South Africa (K417N, E484K, N501Y, D614G) and Brazil (K417N/T, E484K, N501Y, D614G) have been reported that may change the course of the pandemic^6^.

## Results and Discussion

We analyzed whether the newly emerging mutant RBDs may affect the affinity for the viral receptor. In addition, such mutations at the virus-receptor interaction face may alter the ability of RBD-specific antibodies - induced by previous infection or vaccination - to neutralize the mutant viruses. A previous study showed that serum neutralization is not compromised by N501Y (also found in the UK strain B.1.1.7)^7^. In contrast, E484K (found in the South Africa 501Y.V2 and in the Brazilian P.1 strains) was associated with reduced neutralization^8, 9^. Interestingly, studies applying *in vitro* pressure produced similar mutations as those that occurred naturally^10^. To address the questions of antibody binding strength and competition mechanistically, we have expressed the RBD of the Brazilian isolate P.1 exhibiting RBD mutations (K417N, E484K, N501Y), two of which are in the RBM (E484K, N501Y) (Fig. 1A)^11^. Both wild type (RBDwt) and mutant (RBDmut) were purified to homogenicity and the affinity to recombinant ACE2 was determined by Biolayer Interferometry using Octet technology^12^. Fig. 1B shows that the affinity of ACE2 for the novel RBDmut was twice as high as for RBDwt. Having in mind that the affinity of the of SARS-CoV-2 for ACE2 is only 4-fold higher compared to SARS-CoV-1, this factor of 2 is expected to be biologically significant.

**Figure 1.**
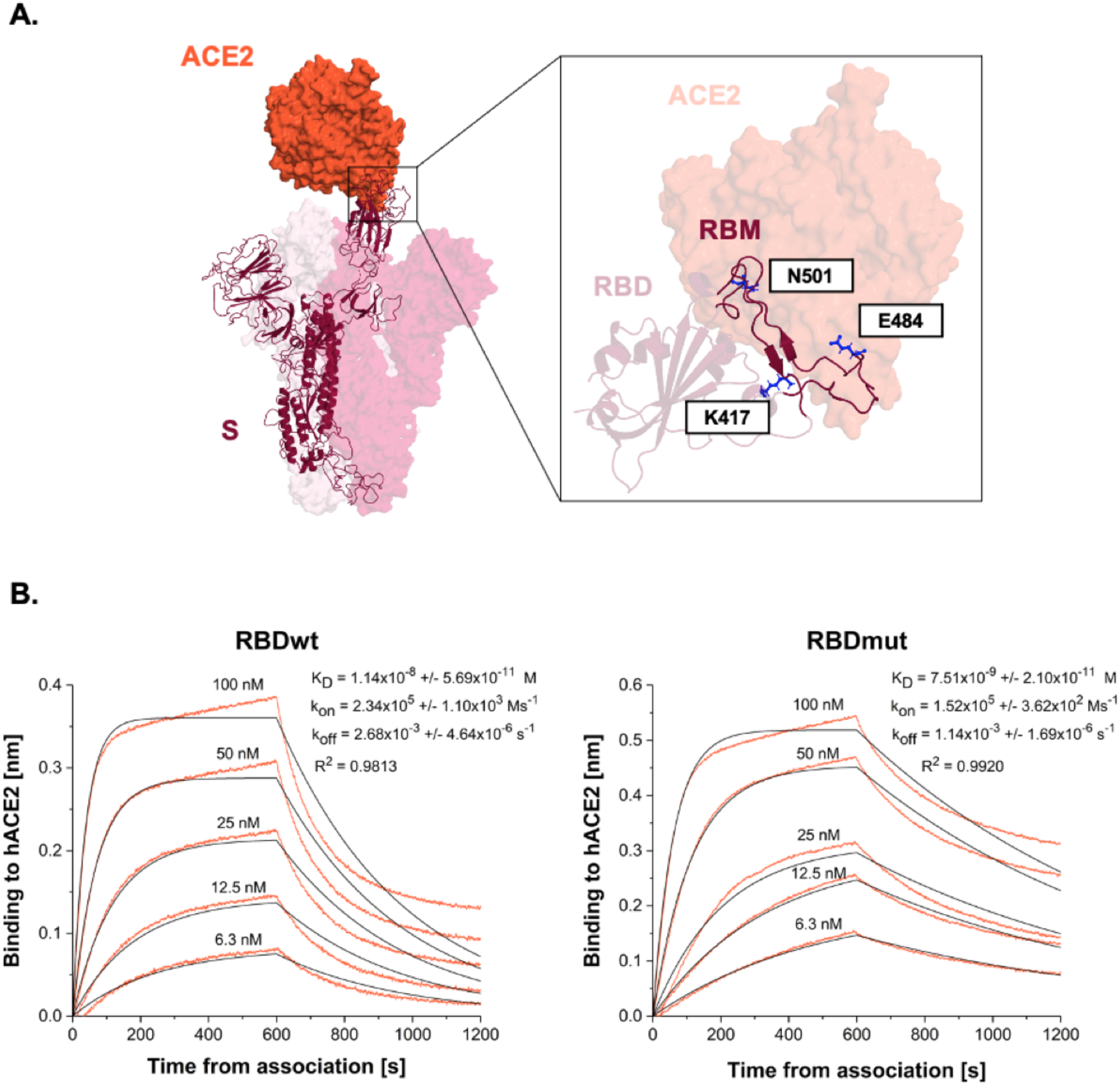
Mutations in the novel Brazilian variant increase affinity of RBD for ACE2. A) The left panel illustrates an S monomer (purple ribbon) bound to hACE2 ectodomain (orange surface). The positions of the key mutations K417N, E484K, and N501K (blue sticks) are indicated on the right panel. B) Binding profiles of RBDwt and RBDmut to ACE2 using biolayer interferometry (BLI).

To determine whether RBDwt-specific immune sera might have a reduced ability to bind to RBDmut we performed ELISA and Biolayer Interferometry. Mice were immunized twice with RBDwt displayed on Virus-Like Particles derived from Cucumber mosaic virus (CuMV_TT_-VLPs) (40 μg, s.c.^13^) and sera were collected 7 days after the second vaccination. Direct binding experiments demonstrated better recognition of RBDwt than RBDmut by murine immune sera (Fig. 2A, B). To assess the capacity of the sera to block the interaction between RBDmut and the viral receptor, we performed competition experiments. The results showed that RBDwt-induced immune sera were clearly inferior at blocking binding of ACE2 to RBDmut, indicating reduced neutralizing capacity of RBDwt specific antibodies for the mutant virus (Fig. 2C). Essentially similar results but with much pronounced effect were obtained for human sera received from convalescent patients. Impaired recognition of RBDmut was paralleled by reduced binding to RBDmut (Fig. 3A, B) and ability to compete with ACE2 binding (Fig. 3C). Thus, these data demonstrate immune escape by RBDmut from polyclonal antibodies induced by vaccination or infection, comparable to what was previously shown with monoclonal antibodies for N501Y^14^ and more importantly for E484K^15, 16, 17^. Even though viral mutations may more strongly affect monoclonal antibodies than sera activity, the latter may also be reduced^18^ as confirmed here. While the studies of monoclonal antibodies are relevant for the characterization of individual epitopes and viral clusters, as well as the potential use of those antibodies to treat COVID-19 patients^19^, the activity of sera is more relevant for immunity as it likely correlates with protection of the increasing number of individuals that have undergone infection and/or vaccination.

**Figure 2.**
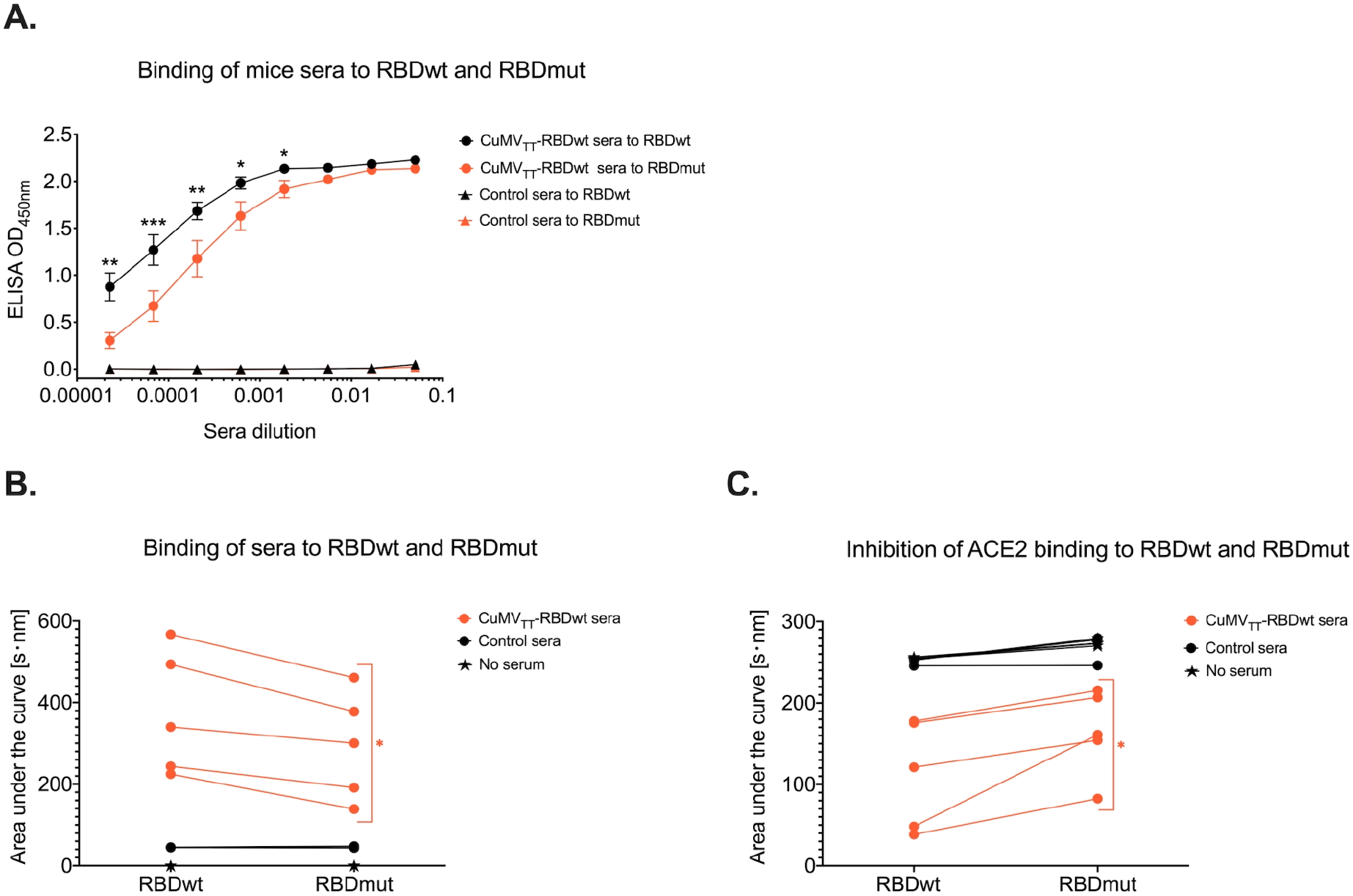
Sera of mice immunized with CuMVTT-RBDwt bind less to RBDmut than RBDwt. A) Binding of sera of 5 CuMVTT-RBDwt immunized mice, and 3 Tris-buffer injected mice, to RBDwt and RBDmut, as determined by ELISA. Direct binding (B) and competitive inhibition of ACE2-mFc interaction to RBDwt and RBDmut (C) by BLI. Sera (dilution 1:20) of the same mice as in A) were used.

**Figure 3.**
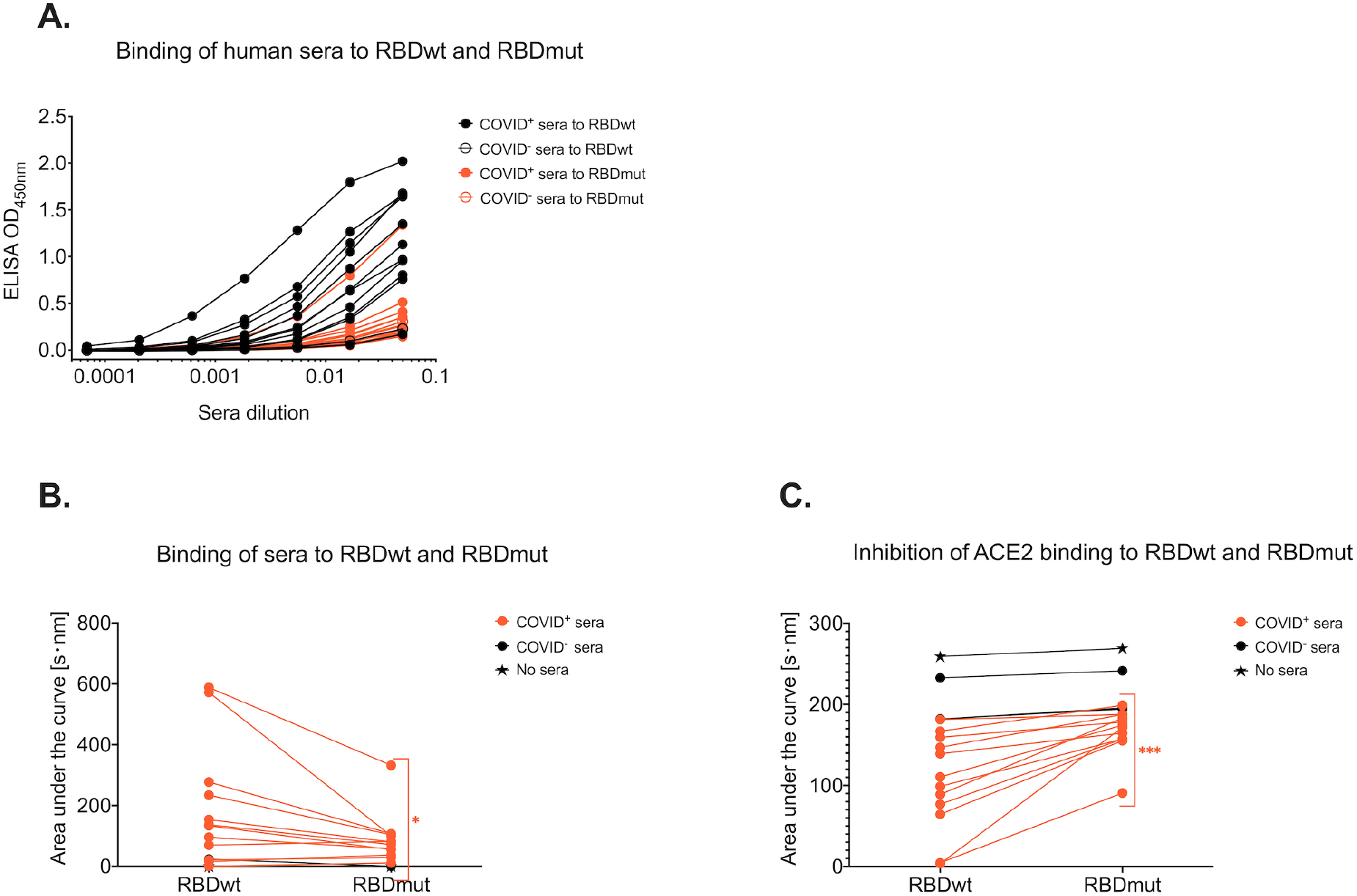
Sera of COVID-19 convalescent patients bind less to RBDmut than RBDwt. A) Binding of sera from 12 COVID-19 convalescent patients and one COVID negative individual to RBDwt and RBDmut, as determined by ELISA. Direct binding (B) and competitive inhibition of ACE2-mFc interaction to RBDwt and to RBDmut (C) were assessed by BLI. Sera (dilution 1:20) of the same individuals as in A) were used.

In summary, our data demonstrate increased affinity of ACE2 for the mutant RBD and reduced ability of RBDwt-induced immune sera to compete with ACE2 binding to RBDmut. These data confirm that the mutant Brazilian SARS-CoV-2 strain P.1 is more infectious^3^ and less susceptible to neutralization by antibodies induced with RBDwt.

## Abbreviations

SARS-CoV-2: Severe Acute Respiratory Syndrome Coronavirus type 2
RBD: Receptor Binding Domain
ACE2: Angiotensin-converting enzyme 2
RBDwt: Receptor Binding Domain wild type
RBDmut: Receptor Binding Domain mutant
COVID-19: Coronavirus disease 2019
BLI: Biolayer interferometry

## Acknowledgements

We thank PD Dr. Alexander Eggel and Dr. Daniel Brigger for providing biotinylated and non-biotinylated ACE2-mFc.

## Methods

### Protein expression and purification

The SARS-CoV-2 receptor-binding domain (RBDwt) and the RBD mutant (RBDmut) (K417N, E484K, N501Y) were expressed using Expi293F cells (Invitrogen, ThermoFisher Scientific, MA, USA). The genes that encode SARS-CoV-2 RBDwt (residues Arg319-Phe541) or RBDmut with a C-terminal 6-His-tag was inserted into pTwist CMV BetaGlobin WPRE Neo vector (Twist Bioscience, San Francisco, USA). The construct plasmids were amplified in *E. coli* XL-1 Blue electrocompetent cells, and were then transfected into Expi293F cells at a density of 3×10^6^ cells/ml using ExpiFectatmine 293 transfection kit (Gibco, ThermoFisher Scientific, MA, USA). The supernatant of cell culture containing the secreted RBDwt or RBDmut was harvested 96 h after infection, and purified by passing His-Trap HP column (GE Healthcare, USA). Collected RBDwt or RBDmut proteins were equilibrated in PBS and kept at −20°C. ACE2-mFc was purchased from Sino Biological (Eschborn, Germany). Biotinylated and non-biotinylated soluble human ACE2 fused to mouse IgG2a Fc proteins were kindly provided by PD Dr. Alexander Eggel (University Clinic of Rheumatology and Immunology, Inselspital, Bern, Switzerland) who received the plasmid from Prof. Peter Kim (Stanford University, CA, USA).

### Generation of the CuMV_TT_-RBDwt vaccine

The RBDwt protein was conjugated to Cucumber mosaic virus - tetanus toxoid (CuMV_TT_) using the linker Succinimidyl 6-(beta-maleimidopropionamido) hexanoate (SMPH) (Thermo Fisher Scientific, Waltham, USA). CuMV_TT_ contains a powerful T cell epitope of the Tetanus toxin (TT) which is capable of delivering strong antibody responses (I). The coupling reactions were performed with mole ratio as following: CuMV_TT_: RBDwt = 1:1, CuMV_TT_: SMPH = 1:7.5. The CuMV_TT_ was firstly linked with SMPH at 25°C, 400 rpm shaking for 30 min, and the RBDwt was reduced by TCEP (Tris (2-carboxyethyl) phosphine) meanwhile. Then the coupling was achieved by mixing the CuMV_TT_ and reduced RBDwt for 3 h at 25°C with 400 rpm shaking. Unreacted SMPH and RBD proteins were removed using Amicon-Ultra 0.5, 100K (Merck-Millipore, Burlington, Mass). Coupling efficiency was calculated by densitometry (as previously described (II)), with a result of approximately 20% to 30%.

### Vaccination of mice

BALB/c mice at the age of 7 weeks were purchased from Envigo (NM Horst, The Netherlands) and kept at SPF animal facility of University of Bern. All animal experiments were performed in accordance with the Cantonal Veterinary guidelines of Bern. Female mice (8-12 weeks old) were immunized by subcutaneous injection with 40 μg CuMV_TT_-RBDwt or Tris buffer (20mM Tris + 5mM EDTA, pH 8.0). The boost of the vaccination was conducted at 24 days after priming. Serum samples were collected 7 days after boost. Five mice were used in CuMV_TT_-RBDwt vaccination group and three mice that were injected with Tris buffer.

### ELISA Assay

Corning half area 96-well plates were coated with 1 μg/ml RBDwt or RBDmut in PBS overnight at 4°C. Then plates were blocked for 2 hours at room temperature with PBS-0.15% Casein. Afterwards, convalescent human sera or immunized mouse sera were added, serially diluted 1:3 and incubated on plates for one hour at room temperature. Bound IgG antibodies were detected with antibody goat anti-mouse IgG-POX antibody (Jackson ImmunoResearch, Cambridgeshire, UK) or goat anti-human IgG-POX antibody (Nordic MUbio, Susteren, The Netherlands). ELISA was developed with tetramethylbenzidine (TMB), stopped by adding equal 1 M H_2_SO_4_ solution, and read at OD450nm.

### RBDwt and RBDmut kinetics by Bio-Layer Interferometry (BLI)

The analysis of binding kinetics of RBDwt and RBDmut to ACE2-mFc was performed by Bio-Layer Interferometry (BLI) using an Octet RED96e (Fortebio) instrument. High precision Streptavidin (SAX, ForteBio, Fremont, CA, USA) biosensors were saturated with 7.5 μg/ml biotinylated ACE2-mFc in BLI assay buffer (PBS, 0.1% BSA, 0.02% Tween 20) for 10 min. RBDwt and RBDmut were prepared as twofold serial dilution (typically 100, 50, 25, 12.5, 6.25 and 3.125 nM) in BLI assay buffer plus buffer blanks. Association was monitored for 10 min and as well dissociation into buffer alone. All steps were performed at 30°C with shaking at 1000 rpm. Kinetic values were calculated by ForteBio data analysis software using a 1:1 binding model.

### BLI-based competitive assay

The ability of the sera of the immunized mice and of the COVID convalescent patient to compete with ACE2 for binding to RBDwt or RBDmut was tested in a sandwich format assay on the Octet RED96e (Fortebio). Anti-penta-His (HIS1K) biosensors were loaded for 10 min with RBDwt His-tag and RBDmut His-tag at a concentration of 7.5 μg/ml in BLI assay buffer. The sensor tips were then dipped in wells containing samples (diluted 1:20 in BLI assay buffer) from convalescent human sera or immunized mouse sera. Association was followed for 10 min. To assess whether the sera can inhibit the binding of ACE2 to RBDwt or RBDmut tips were then placed in wells containing ACE2-mFc at a concentration of 50 nM and association was measured for 10 min. For control two additional tips with BLI buffer were used, one for baseline and one without serum sample to determine binding of ACE2-mFc alone. The results are expressed of individual mice. The response data were normalized using ForteBio data analysis software version1.2.0.1.55.

### Data and statistical analysis

All statistical tests were performed using GraphPad PRISM 6.0 (GraphPad Software, Inc.). ELISA data in graphs are displayed as OD_450_ values of individual mice ± SEM. BLI data are shown as area under the curve for each individual mice or humans. Comparison between RBDwt and RBDmut were analyzed by paired two-tailed Student‘s t-test. α=0.05 and statistical significance are displayed as p≤0.05 (*), p≤0.01 (**), p≤0.001 (***).

